# Detection of a reassortant swine- and human-origin H3N2 influenza A virus in farmed mink in British Columbia, Canada

**DOI:** 10.1101/2024.05.27.596080

**Authors:** Kevin S. Kuchinski, John Tyson, Tracy Lee, Susan Detmer, Yohannes Berhane, Theresa Burns, Natalie A. Prystajecky, Chelsea G. Himsworth

## Abstract

In December 2021, influenza A viruses (IAV) were detected in a population of farmed mink in British Columbia, Canada. Based on genomic sequencing and phylogenetic analysis, these IAVs were subtyped as H3N2s that originated from reassortment of swine H3N2 (clade 1990.4h), human seasonal H1N1 (pdm09), and swine H1N2 (clade 1A.1.1.3). This reassortant has been subsequently observed in swine in several Midwest American states, as well as in swine and turkeys in Ontario, suggesting its spillover into farmed mink in British Columbia was incidental to its broader dissemination in North American swine populations. These detections reaffirm the need for extensive genomic surveillance of IAVs in swine populations to monitor reassortments that might become public health concerns. They also highlight the need for closer surveillance of IAVs in mink to preserve animal health, protect agricultural interests, and monitor potential zoonotic threats.

## Introduction

Influenza A viruses (IAVs) infect diverse mammalian hosts due to conserved glycosylation of cell surface proteins that act as viral receptors. Specifically, sialic acids linked to galactose by α2,6-linkages (SA α2,6 GAL) within respiratory tract epithelium are exploited by IAVs for cell attachment and entry in humans and other mammals like swine and mustelids^1,2^. This has made swine a common source of zoonotic influenzas, and it has made mustelids, especially ferrets, a popular animal model for studying IAV respiratory infections^2–4^.

Swine and certain mustelids, notably mink, also have sialic acid receptors with α2,6 and α2,3 galactose linkages (SA α2,3 GAL) on their intestinal epithelium, the latter of which is the preferred receptor for IAVs of avian origin^2,5^. Consequently, swine have been recognized as “mixing vessels” for avian- and mammalian-origin IAVs, implicating them in the emergence of past pandemic IAV strains^6,7^. Increasingly, a similar mixing vessel role has been proposed for farmed mink, with suggestions of increased risk due to the high degree of similarity between human and mustelid respiratory physiology^2^. Indeed, mink can be infected, both naturally and experimentally, with IAVs originating from birds, swine, and humans^8–11^. These general concerns about the role of mink in zoonotic disease were reinforced during the COVID-19 pandemic by widespread spillover of SARS-CoV-2 into farmed mink populations, with worries of potential spillback of novel variants into humans^12,13^. This resulted in unprecedented surveillance and molecular testing of farmed mink populations with respiratory disease.

IAVs infections in mink, however, are frequently asymptomatic and/or manifest in disease only when there is co-infection with another viral or bacterial pathogen, therefore it is likely that cases and outbreaks are under detected^14,15^. The source of IAV infections in farmed mink is also unclear. It has been hypothesized that contamination of the mink farm environment by wild birds could be source of infection, as could contact with infected farm workers^10,11,16^. Most commonly, infection is suspected to result from feeding raw, infected pork and poultry products^9,16–18^. But overall, relatively little is known about the incidence, origin, or significance of IAV infections in farmed mink.

In December 2021, IAV infections were incidentally detected in a population of farmed mink in the province of British Columbia (BC), Canada during a SARS-CoV-2 outbreak investigation and surveillance program. In this study, we performed genomic characterization of the IAV associated with this outbreak and investigated its source.

## Materials and Methods

### Specimen collection

Oropharyngeal specimens were collected from manually restrained mink by inserting a sterile polyester swab into the mink’s mouth through a gag (either a 3cc syringe tube or a short length of sterilized 1 inch diameter copper pipe). The swab was swirled several times as far back in the oral cavity as possible, then placed into viral transport medium for storage and shipment.

### Nucleic acid extraction and diagnostic testing

Total nucleic acids were extracted from 200 μL of inoculated viral transport medium using the MagMax™ Express 96 Nucleic Acid Extractor (Applied Biosystems) and the MagMax Viral/Pathogen Nucleic Acid Isolation Kit (Thermo Fisher Scientific) according to the manufacturer’s recommendations. Extracted nucleic acids were assayed by RT-qPCR for SARS-CoV-2, IAV, influenza B virus, and respiratory syncytial virus as previously described^19^.

### Genome sequencing and bioinformatic analysis

Influenza A virus cDNA was generated from specimen RNA using the amplification method described by Zhou *et al.* with modifications to the primers to improve amplicon complexity and specificity^20^. Sequencing libraries were constructed from cDNA using the Illumina DNA Prep Kit following a modified condensed protocol^21^. Libraries were sequenced in-house on an Illumina MiSeq platform at the BC Centre for Disease Control Public Health Laboratory (PHL) using a V2 300 cycle micro kit (MS-103-1002).

Influenza A virus genome segment sequences were generated using FluViewer (v0.1.9) (https://github.com/KevinKuchinski/FluViewer). Default parameters were used, except for minimum read depth for base calling (-D), which was set to 10. Complete segment sequences (>= 95% segment coverage and >= 95% non-ambiguous base calls) were remotely queried against the NCBI Nucleotide Collection on 19 Feb 2024 using blastn (v2.14.1+) with subjects limited to the influenza A virus taxon (txid 11320)^22^. Alignments from blastn were grouped by subject sequence, then the median bitscore was calculated for each subject sequence to measure the similarity of each IAV reference sequence to the IAVs from the mink outbreak collectively. Median bitscores were used to identify the best-matching reference sequences for each genome segment. This provided a list of IAVs in GenBank with at least one segment closely related the IAVs from the mink outbreak. GenBank records for these closely related IAV strains were manually inspected, and a sample of them was selected for phylogenetic analysis based on the availability and completeness of all 8 genome segments. Multiple sequence alignments were generated with MUSCLE (v5.1.linux64), which were then used to build maximum likelihood phylogenetic trees using PhyML (v3.3.20220408)^23,24^. Each tree was constructed with 100 bootstrap replicates. Clades for H1 and H3 swine viruses were determined using the subspecies classification tools for Orthomyxoviridae provided by the Bacterial and Viral Bioinformatics Resource Centre (https://www.bv-brc.org).

## Results

On 2 May 2021, an outbreak of SARS-CoV-2 was detected on an American mink farm (*Neovison vison*) in BC through a passive surveillance program. This program had been instituted by the Ministry of Agriculture and Food (MAF) in response to two previous SARS-CoV-2 outbreaks on mink farms in BC. The outbreak continued until April 2022, at which point the farm was depopulated. No unusual clinical signs were reported in the herd, nor was there increased incidence of respiratory disease or mortality. During the surveillance period, SARS-CoV-2 prevalence and evolution was monitored by testing a random stratified sample of 65 mink every two weeks. Testing was conducted by the BC Centre for Disease Control Public Health Laboratory (PHL), where SARS-CoV-2 assays had been multiplexed with IAV, influenza B virus, and respiratory syncytial virus for higher laboratory throughput during the COVID-19 pandemic. This led to the incidental detection of IAV in 17 of 65 mink specimens on 3 Dec 2021.

To confirm this diagnosis and further characterize these IAVs, whole genome sequencing was conducted. Fifteen complete HA segment sequences and 16 complete NA segment sequences were obtained. When queried against the NCBI Nucleotide Collection (Table S1), these HA and NA sequences had their best alignments to swine-origin H3 and N2 sequences. We also obtained 78 complete internal segment sequences from these mink specimens (9 PB2, 8 PB1, 12 PA, 16 NP, 17 M, and 16 NS). These internal segments had their best alignments to a mixture of swine-origin and human seasonal IAVs (Table S1), suggesting the H3N2 viruses detected in these mink originated through reassortment.

To investigate this apparent reassortment, we constructed phylogenetic trees for each segment using sequences from the mink specimens, closely related reference viruses from NCBI, and unpublished sequences from Canadian swine (Figures S1-S8). To assess possible epidemiological connections, these trees included four H3N2 viruses detected on local BC swine farms over the preceding six years (Figures S1-S8, pink leaves). The phylogenies suggested that the PA, HA, NA, M, and NS segments of the mink viruses were derived from an H3N2 lineage that has been observed in Saskatchewan swine since 2017 (Figures S1-S8, purple leaves). This progenitor lineage belonged to clade 1990.4, also known as Cluster IV, the most prevalent swine H3N2 in North America in recent years^25–27^. The trees also indicated that this H3N2 progenitor was itself a reassortant of earlier swine H3N2 (clade 1990.4) and H1N2 (clade 1A.1.1.3) viruses circulating in Western Canadian in the preceding years (Figures S1-S8, red and blue leaves respectively).

Phylogenetic analysis also indicated that the PB2, PB1, and NP segments of these mink viruses were derived from human seasonal H1N1pdm09 (Figures S1-S8, green leaves). Their best-matching human seasonal H1N1pdm09s circulated in 2018/19, suggesting reassortment of these segments had occurred 2 to 3 years prior to the mink farm outbreak. Temporal separation between reassortment and infection of these mink was further supported by the comparatively long branches separating mink-origin sequences from the human-origin sequences with which they formed clades.

Figure 1 summarizes the proposed series of reassortments that gave rise to the H3N2 lineage detected in this mink farm outbreak. This IAV’s constellation of genome segments has been subsequently observed in swine in Iowa and Minnesota in 2022 as well as turkeys and swine in Ontario in 2022 and 2023 (Figures S1-S8, orange leaves). This pairing of HA and NA segments was also observed in swine in Missouri in 2021 (Figures S4 and S6, orange leaves), but it is unknown if this was an additional instance of this full genome constellation or a subsequent reassortant because internal segment sequences were not submitted for the IAVs from Missouri. Taken together, these results suggest that the H3N2 lineage detected in the mink outbreak has become widespread in North American swine, and infection of these mink with this lineage was incidental to its broader dissemination in swine.

**Figure 1:**
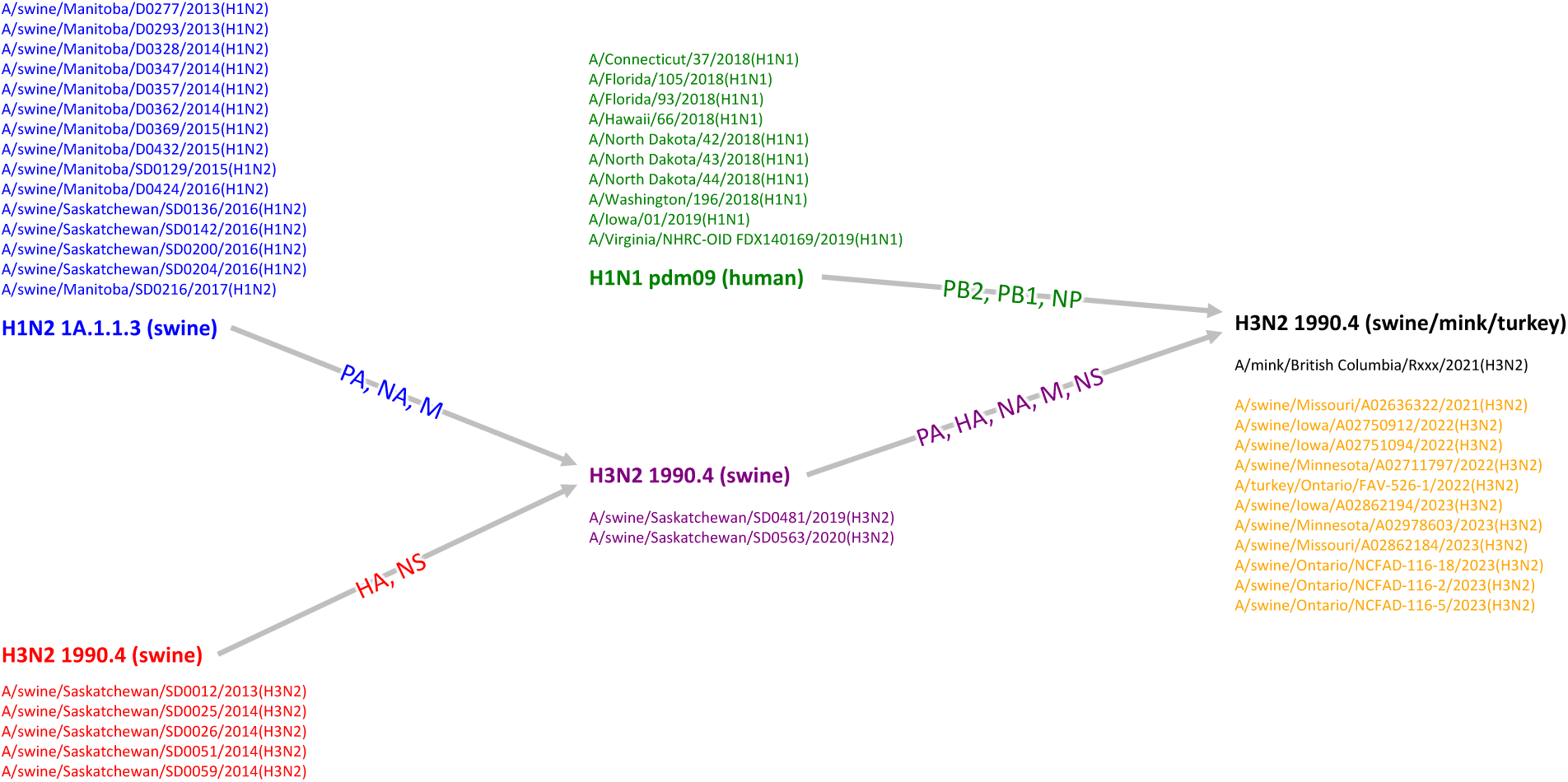
Simplified schematic of re-assortments resulting in the H3N2 influenza A virus (IAV) detected in farmed mink. The origin of each genome segment in the mink H3N2 virus (black) was determined based on their best-matching reference sequences and the phylogenies presented in Figures S1-S8. The PB2, PB1, and NP segments were derived from human season H1N1 pdm09 *c*. 2018/19 (green). The PA, HA, NA, M, and NS segments were derived from a swine H3N2 clade 1990.4 virus (purple), which itself was a reassortant of an earlier swine H3N2 clade 1990.4 (red) and a swine H1N2 clade 1A.1.1.3 (blue). This reassortant that was detected in mink has been subsequently detected in swine and turkeys in Ontario and American Midwest states (orange).

We interviewed the mink producers and their primary care veterinarians to assess several scenarios by which these minks could have been exposed to swine-origin IAVs. First, we ruled out direct contact between mink and infected swine because no swine were located on the mink farm premises. We also found no evidence of transmission via fomites; mink farm personnel could not recall having any contact with swine or visiting any premises with swine in the months preceding the outbreak. Exposure via contaminated pork was also considered, but the producers did not recall feeding the mink any pork products.

Swine-origin H3N2 infections have been reported in turkeys^28^, so we also investigated potential direct and indirect exposures to turkeys. As with swine, there were no turkeys on the mink farm premises and no personnel or equipment had been shared with turkey farms. Mink had been fed raw poultry necks and backs obtained from a local processing plant, but it was not known if the necks and backs were from turkeys or other poultry species.

We also considered if the mink had been exposed to humans or wild birds infected with swine-origin IAVs. There were no reported cases of humans infected with swine-origin IAVs in BC in 2021, and no farm personnel could recall any influenza-like illness around the time of the outbreak. Exposure to birds infected with swine-origin IAVs seemed equally unlikely; no swine-origin IAVs were detected through avian influenza surveillance programs during the 2021/22 season. This included passive surveillance of wild birds as well as novel environmental surveillance of avian habitats using targeted genomic sequencing of sediment from local wetlands^29^. Several wetlands in the same region as the mink farm were monitored, resulting in detections of diverse IAVs, but none were swine-origin. Taken together, this indicated that the mink were probably not exposed to humans or wild birds infected with swine-origin IAVs because swine-origin IAVs were not circulating in the human population or local wild bird community.

## Discussion

Our investigation could not conclude how these mink became exposed to swine-origin IAVs. Additional modes of transmission were considered, but they could not be assessed due to lack of available data. For instance, wild mustelids have been reported to visit mink farms and interact with captive animals resulting in the transmission of viruses^30^; it is possible that wild mustelids may have visited this farm unnoticed after becoming infected with IAVs on another premise where swine are raised.

We also considered environmental transmission of IAVs between local swine and poultry operations and this mink farm. Shortly before the outbreak, on 14 November 2021, a severe atmospheric river system inundated the region where the mink farm is located,^31^ resulting in a regional state of emergency and historic flooding that submerged parts of the mink farm. It is conceivable that swine or poultry excreta were transported onto the premises in flood waters. Transmission of IAVs between birds in aquatic environments is well-established, and long-term persistence of IAVs in these aquatic environments has been demonstrated^32,33^. Water-borne transmission of IAVs into mammalian livestock has also been proposed in previous outbreaks^34^.

Another potential form of environmental transmission is atmospheric dispersion through wind. Airborne transmission has been invoked to explain spread between farms in both poultry and swine IAV outbreaks, and this has been corroborated to varying degrees with epidemiologic modeling, detection of viral RNA in air specimens, and IAV isolation from air specimens by viral culture^35–39^. Mathematical modeling has proposed wind-borne transmission might be possible over distances of up to 25 kilometers^38^. We did not find any reports in the literature of IAV detection in air specimens further than 2.1 kilometers downwind from infected barns, however, and these were weak detections of viral RNA without successful virus isolation^35^. Two swine farms were located within 10 km of the site of the mink outbreak; one 2.2 kilometers away and the other 9.0 km away. Eleven turkey farms were located within 10 km of the mink farm.

Ultimately, the limited extent of genomic surveillance for IAVs in local swine and poultry populations constrained our ability to identify a local source for the outbreak. It also restricted our ability to assess the plausibility of different transmission routes. Although IAV is a reportable disease in swine and poultry in BC, the passive nature of surveillance programs combined with the potential for asymptomatic or unremarkable infections means that under-reporting and under-detection is likely. Indeed, only 4 contemporaneous, local swine-origin H3N2 IAV genomes were available for analysis, opportunistically detected through an unrelated research study, and these viruses were not related to the mink farm outbreak. This suggests that IAV diversity within swine populations is under-characterized. This was further indicated by limited detections of IAVs with the same genome constellation as far afield as Iowa, Minnesota, Missouri, and Ontario. This suggests that this IAV reassortant was able to disseminate across North America largely unnoticed. The uncomfortable corollary is that many other reassortant IAVs are likely emerging and disseminating unobserved within large, transnational, commercial swine populations.

Similarly, this study suggests that IAV spillovers into mink may be underestimated. After all, these detections were incidental and relied on: 1) an historic SARS-CoV-2 global pandemic that led to unprecedented molecular testing of mink with respiratory signs, and 2) the fortuitous multiplexing of SARS-CoV-2 with IAV in the laboratory where the mink specimens were tested. Without these circumstances, this outbreak would likely have passed unrecognized, suggesting IAV spillovers into mink are more common than assumed. Indeed, serological evidence for frequent spillovers of diverse avian, swine, and human seasonal IAVs into mink has been previously reported^8–11,16–18,40,41^. For example, a survey of sera collected from mink slaughterhouses in Eastern China between 2016 and 2019 revealed 47.3% and 11.4% seroconversion to H1N1 and H3N2 IAVs, which were presumed to be human seasonal lineages due to an association with incidences of these subtypes in local human populations^41^. Furthermore, surveys of sera from mink slaughterhouses and live mink herds between 2013 and 2019, also in Eastern China, showed up to 47.5% seroconversion to avian-origin H9N2, the most common subtype infecting Chinese poultry at the time^17,41,42^.

Spillover of diverse IAVs into mink raise several biosecurity and infection prevention and control issues. For mink producers, primary care veterinarians, and agriculture officials, there are questions of animal welfare and economic impact. It has been observed that mink infections with IAVs do not typically produce remarkable disease^9,14,15^, which was indeed the case in this outbreak. This does not preclude severe disease in mink after future IAV spillovers, however. Indeed, natural infections of mink with human season H1N1 and avian-origin H5N1 viruses have been shown to induce severe disease, especially in kits^10,16,43^. It also possible that IAV impacts on mink are underestimated and largely unattributed due to limited molecular testing and confirmatory genomics for mink with respiratory disease.

These spillovers also raise issues for public health officials. Similarities between human and mustelid respiratory physiology suggest overlapping susceptibility to certain IAV lineages, as evidenced by the apparently frequent spillover of human seasonal IAVs into mink noted above. This means that IAVs detected in mink must be assessed as potential spillover threats to humans. Furthermore, the detection of avian- and swine-origin lineages circulating in mink raises the possibility that these animals may facilitate IAV genome segment re-assortment, similar to the mixing vessel role played by swine^2,44^. High seroconversion rates also suggest that mink have a larger exposure interface with poultry and swine populations than humans, which may allow them to serve as a selective conduit into human populations for reassortants arising in other livestock.

While this creates spillover risk, it also creates surveillance opportunities. The mink industry could be a valuable sentinel for reassortant zoonotic IAVs, especially those with high potential to successful replicate within and transmit between human hosts. Fortunately, no spillover between mink and humans was observed in this outbreak, and these IAVs were assessed to pose no immediate threat to public health. Furthermore, any hypothetical threat from this particular IAV lineage via mink would appear trivial compared to the threat from the much vaster transnational commercial swine populations in which this lineage is apparently widespread.

While this study was unable to identify the local source of this outbreak, it did provide valuable insights. Foremost, it highlighted the need for more extensive genomic surveillance of IAVs in livestock, especially swine and mink. This would assist outbreak investigations and further reveal the type of exposures that are responsible for IAV infections in mink. Drawing these connections may also advance our understanding of environmental transmission between farms, which would improve infection prevention and control practices for all livestock industries impacted by IAV spillovers. Finally, heightened genomic surveillance of IAVs in swine and mink would increase detections of well-adapted reassortments, improving pandemic preparedness.

## Acknowledgements

We are deeply grateful to Terry Engebretson and Dr. Dave McDermid for their invaluable contributions to this project. We also thank Dr. Agatha Jassem, Frankie Tsang, and the virology program at the BC CDC PHL for their assistance with molecular testing of mink specimens.

## Data Availability

IAV genome segment sequences have been uploaded to the Global Initiative for Sharing All Influenza Data (GISAID) EpiFlu database under the accession numbers EPI_ISL_19132429 and EPI_ISL_19166148 to -63.

---

**Figure S1:**
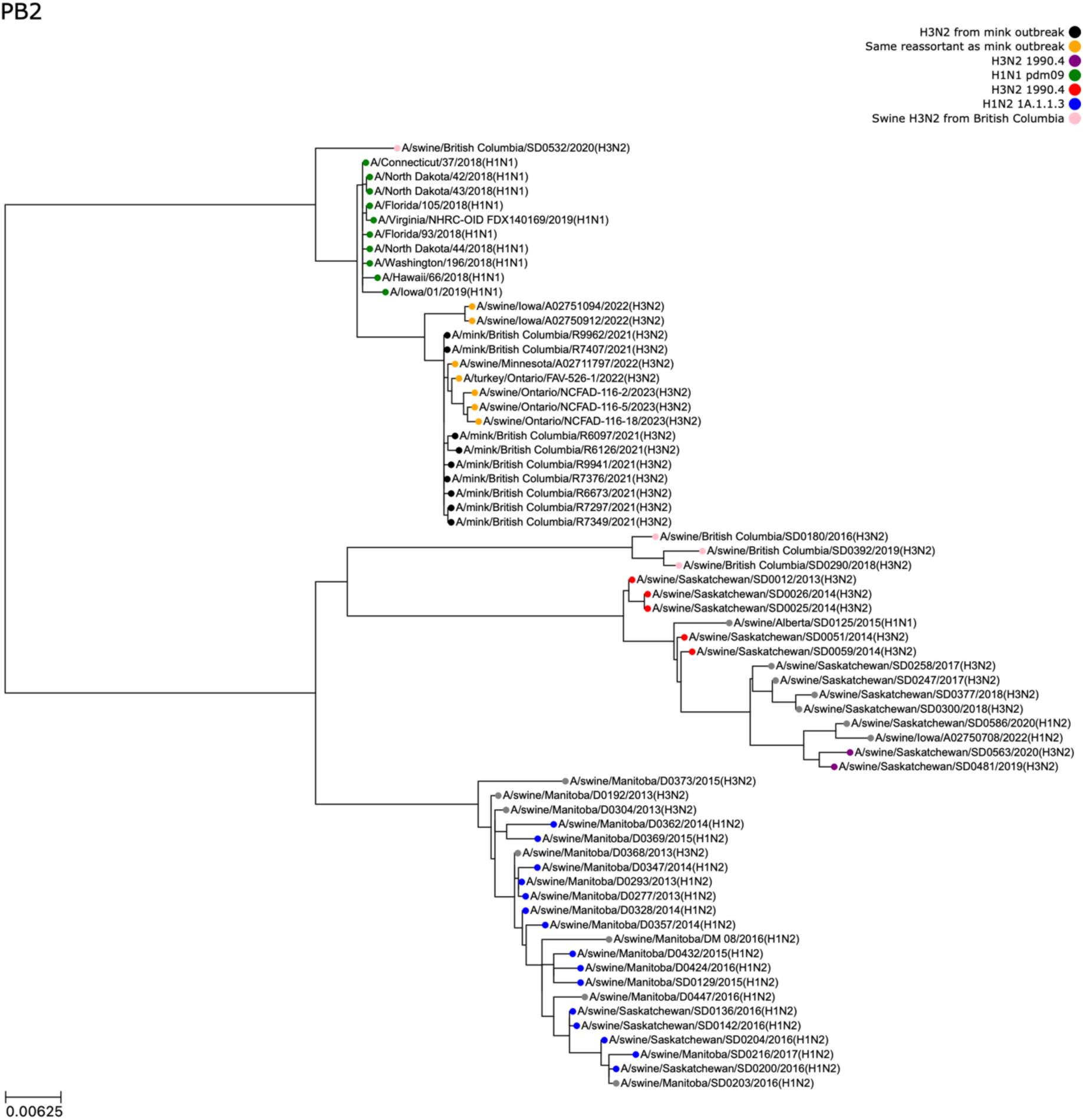
Phylogenetic tree depicting evolutionary relationships between PB2 segments from the influenza A virus (IAV) detected in mink outbreak and other closely-related IAVs. Other IAV sequences were obtained from NCBI or the Canadian Food Inspection Agency. PB2 segments from reference viruses were included in the tree if any segment from those viruses was one of the top matches for any segment of the mink virus. Phylogeny indicates the IAV detected in the mink outbreak (black leaves) obtained its PB2 segment from human seasonal H1N1 pdm09 (green leaves) *c.* 2018/19.

**Figure S2:**
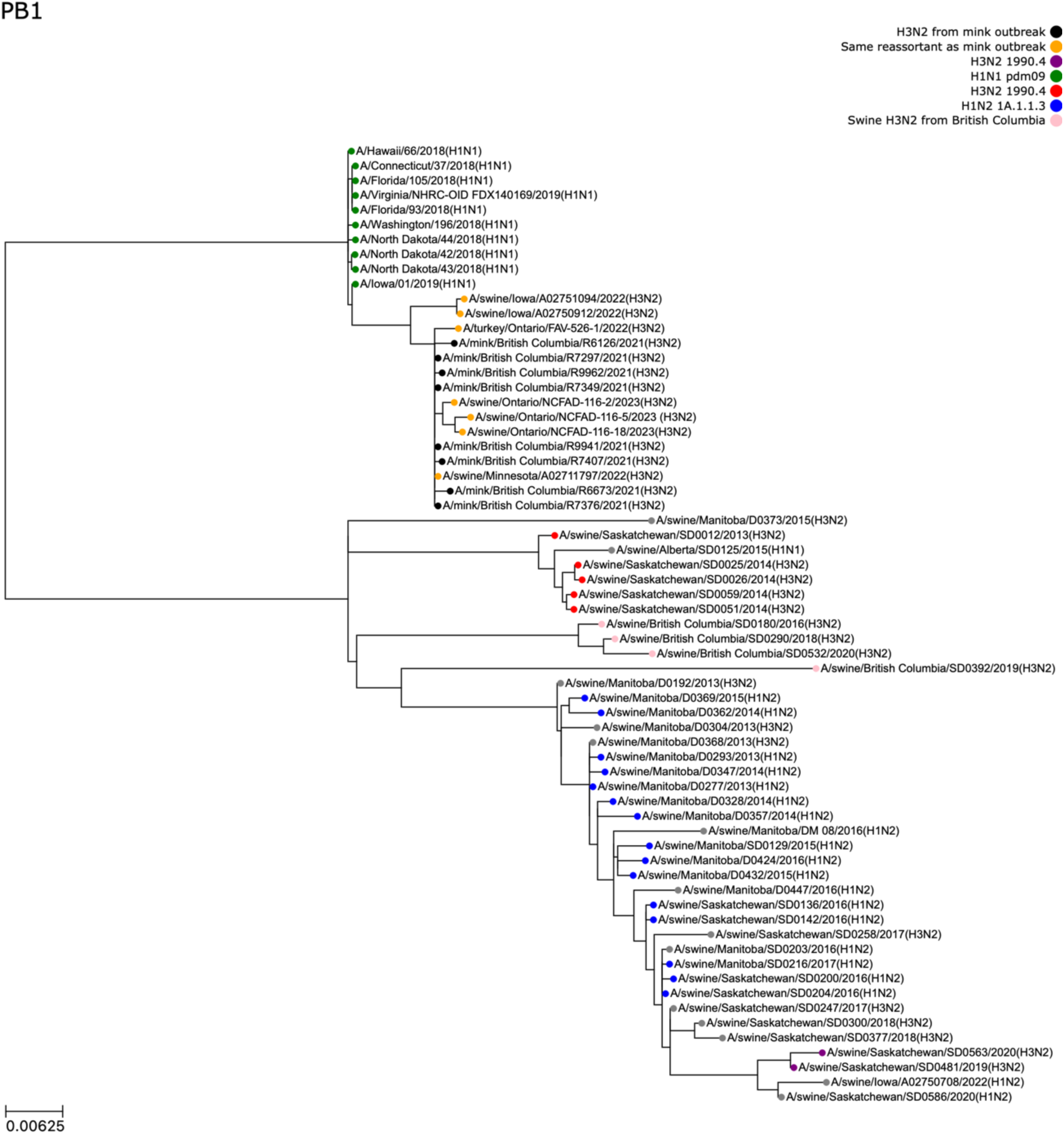
Phylogenetic tree depicting evolutionary relationships between PB1 segments from the influenza A virus (IAV) detected in mink outbreak and other closely-related IAVs. Other IAV sequences were obtained from NCBI or the Canadian Food Inspection Agency. PB1 segments from reference viruses were included in the tree if any segment from those viruses was one of the top matches for any segment of the mink virus. The phylogeny indicates the IAV detected in the mink outbreak (black leaves) obtained its PB1 segment from human seasonal H1N1 pdm09 (green leaves) *c.* 2018/19.

**Figure S3:**
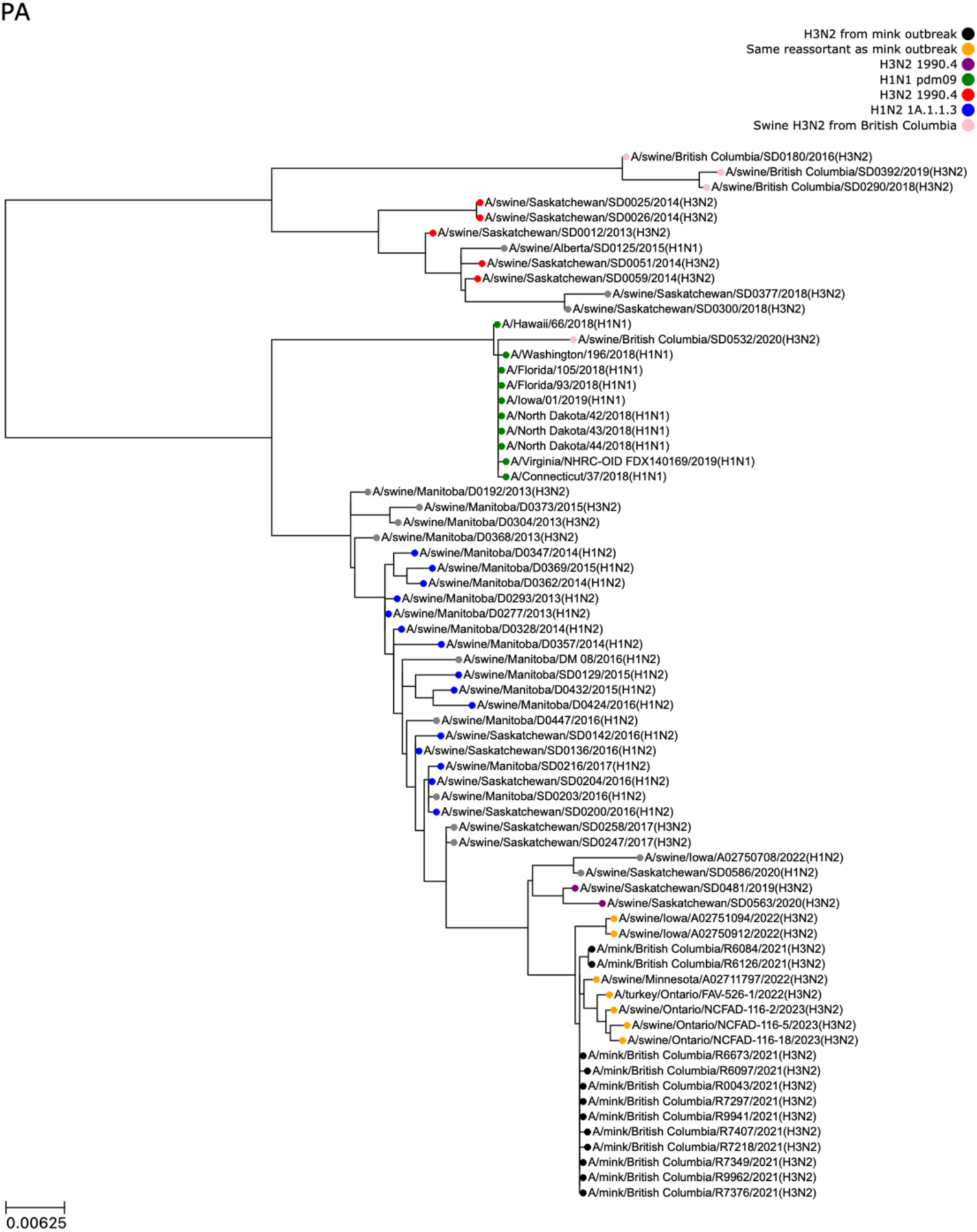
Phylogenetic tree depicting evolutionary relationships between PA segments from the influenza A virus (IAV) detected in mink outbreak and other closely-related IAVs. Other IAV sequences were obtained from NCBI or the Canadian Food Inspection Agency. PA segments from reference viruses were included in the tree if any segment from those viruses was one of the top matches for any segment of the mink virus. The phylogeny indicates the IAV detected in the mink outbreak (black leaves) obtained its PA segment from a swine H3N2 clade 1990.4 virus (purple leaves), which itself obtained this PA segment from a swine H1N2 clade 1A.1.1.3 virus (blue leaves).

**Figure S4:**
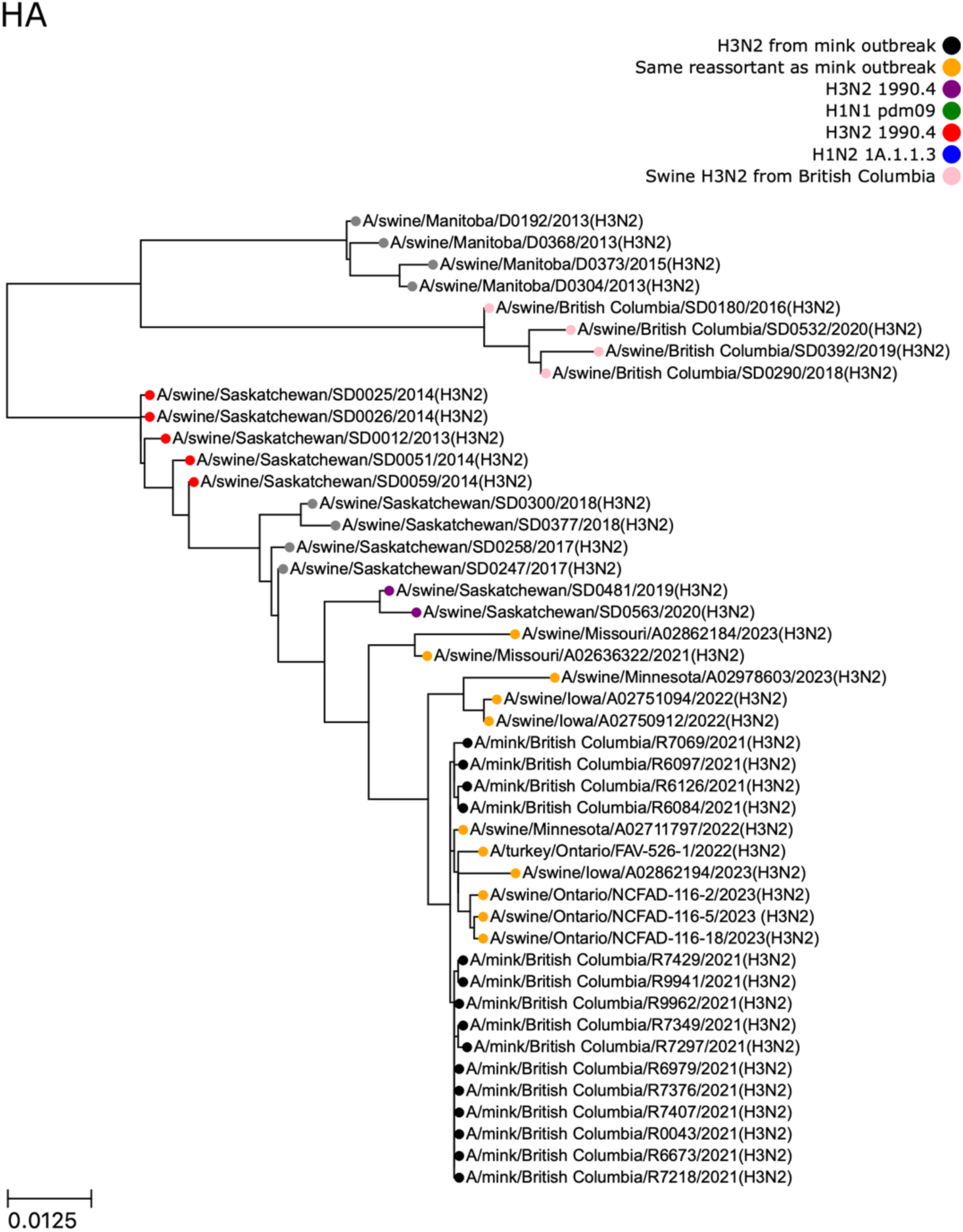
Phylogenetic tree depicting evolutionary relationships between HA segments from the influenza A virus (IAV) detected in mink outbreak and other closely-related IAVs. Other IAV sequences were obtained from NCBI or the Canadian Food Inspection Agency. HA segments from reference viruses were included in the tree if any segment from those viruses was one of the top matches for any segment of the mink virus. The phylogeny indicates the IAV detected in the mink outbreak (black leaves) obtained its HA segment from a swine H3N2 clade 1990.4 virus (purple leaves), which itself obtained this HA segment from an earlier H3N2 clade 1990.4 virus (red leaves).

**Figure S5:**
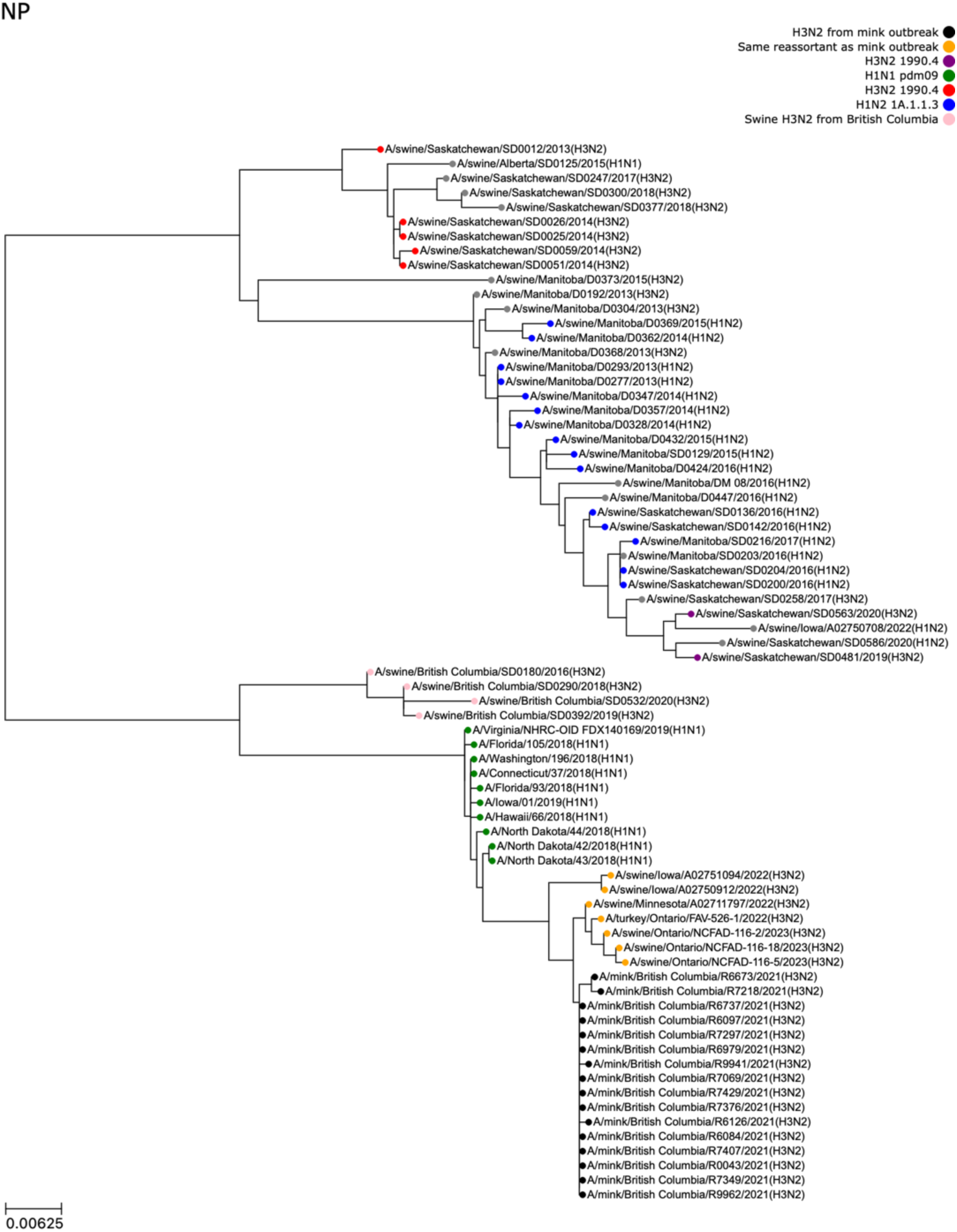
Phylogenetic tree depicting evolutionary relationships between NP segments from the influenza A virus (IAV) detected in mink outbreak and other closely-related IAVs. Other IAV sequences were obtained from NCBI or the Canadian Food Inspection Agency. NP segments from reference viruses were included in the tree if any segment from those viruses was one of the top matches for any segment of the mink virus. The phylogeny indicates the IAV detected in the mink outbreak (black leaves) obtained its NP segment from human seasonal H1N1 pdm09 (green leaves) *c.* 2018/19.

**Figure S6:**
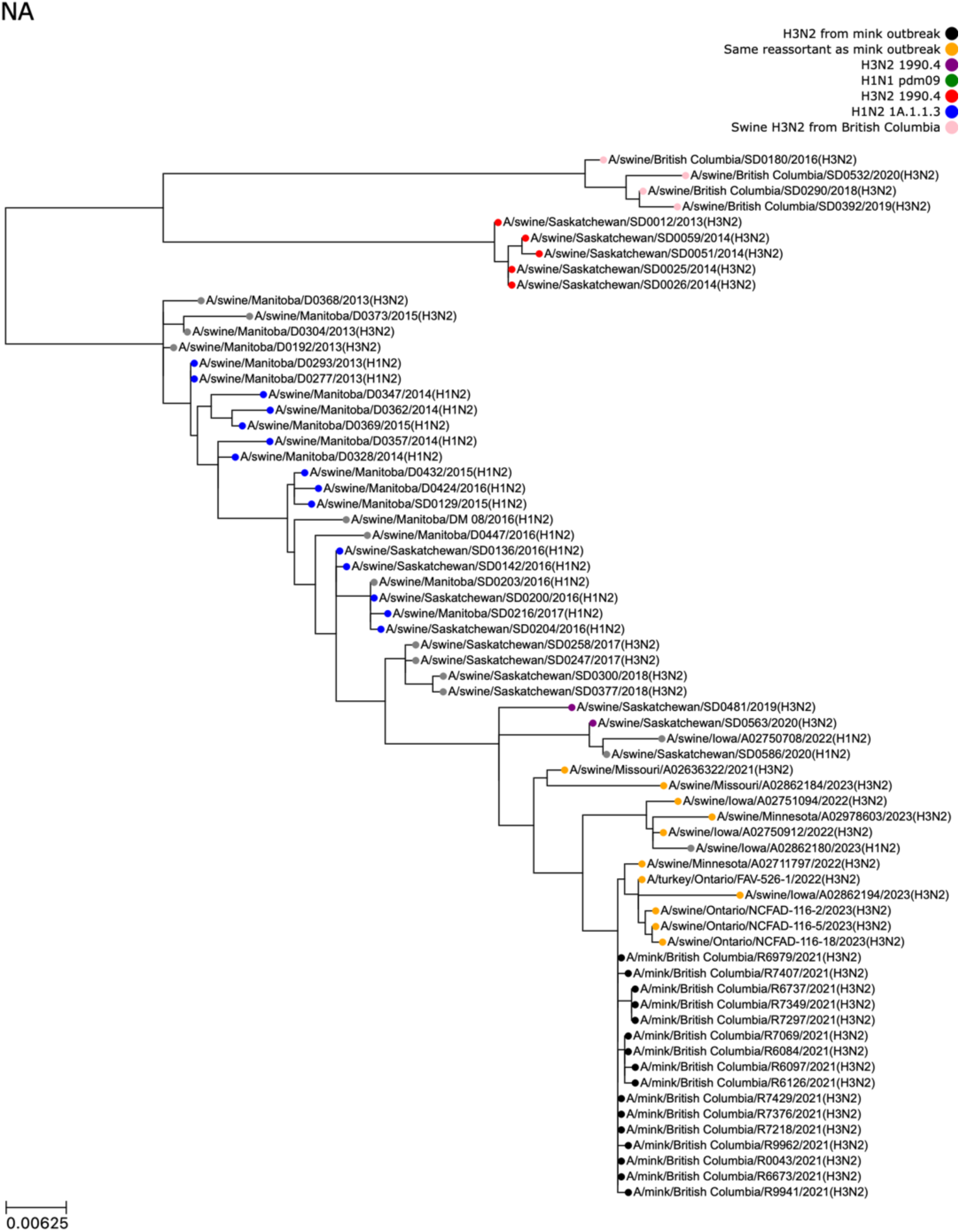
Phylogenetic tree depicting evolutionary relationships between NA segments from the influenza A virus (IAV) detected in mink outbreak and other closely-related IAVs. Other IAV sequences were obtained from NCBI or the Canadian Food Inspection Agency. NA segments from reference viruses were included in the tree if any segment from those viruses was one of the top matches for any segment of the mink virus. The phylogeny indicates the IAV detected in the mink outbreak (black leaves) obtained its NA segment from a swine H3N2 clade 1990.4 virus (purple leaves), which itself obtained this NA segment from an earlier H1N2 clade 1A.1.1.3 virus (blue leaves).

**Figure S7:**
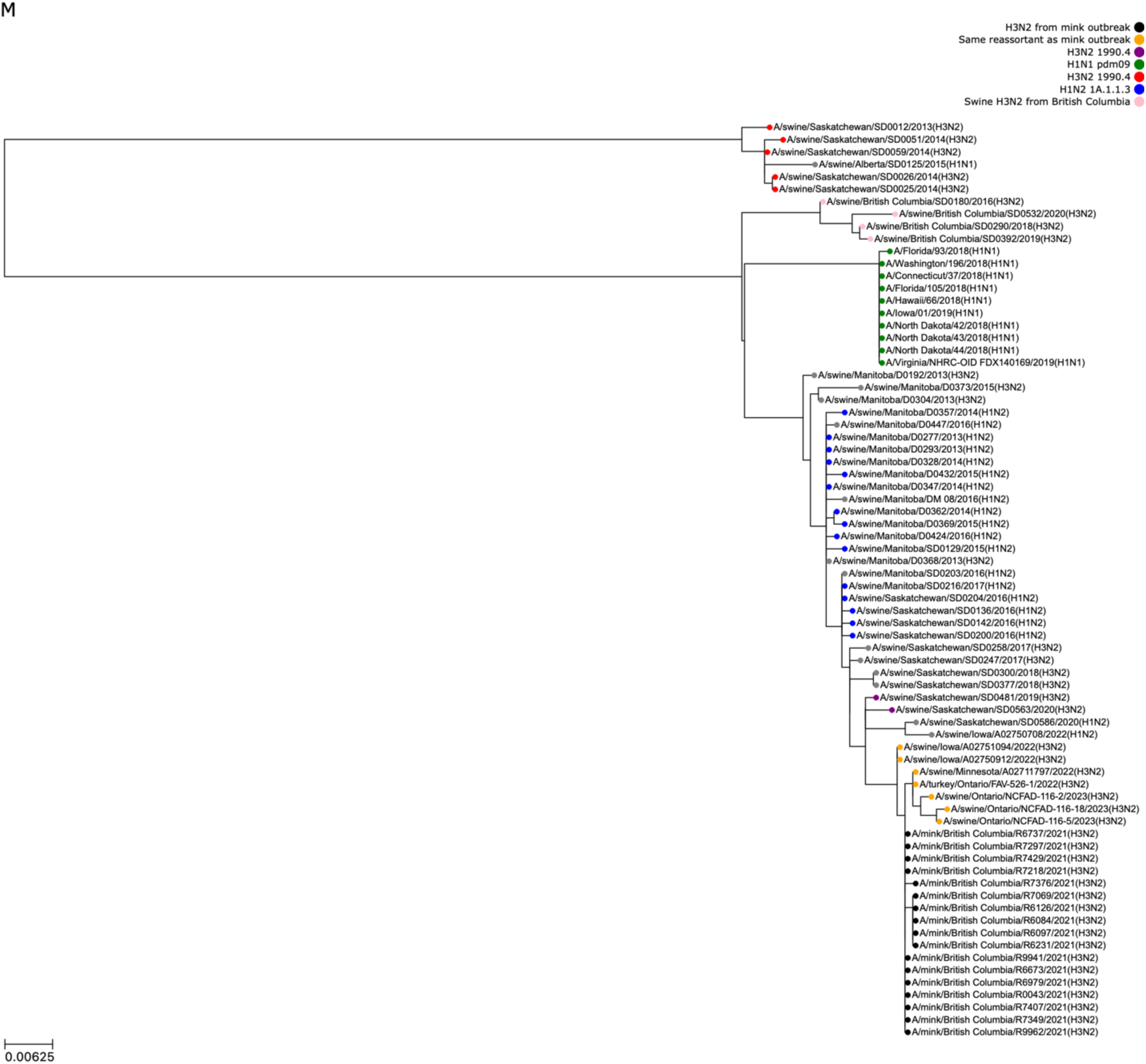
Phylogenetic tree depicting evolutionary relationships between M segments from the influenza A virus (IAV) detected in mink outbreak and other closely-related IAVs. Other IAV sequences were obtained from NCBI or the Canadian Food Inspection Agency. M segments from reference viruses were included in the tree if any segment from those viruses was one of the top matches for any segment of the mink virus. The phylogeny indicates the IAV detected in the mink outbreak (black leaves) obtained its M segment from a swine H3N2 clade 1990.4 virus (purple leaves), which itself obtained this M segment from an earlier H1N2 clade 1A.1.1.3 virus (blue leaves).

**Figure S8:**
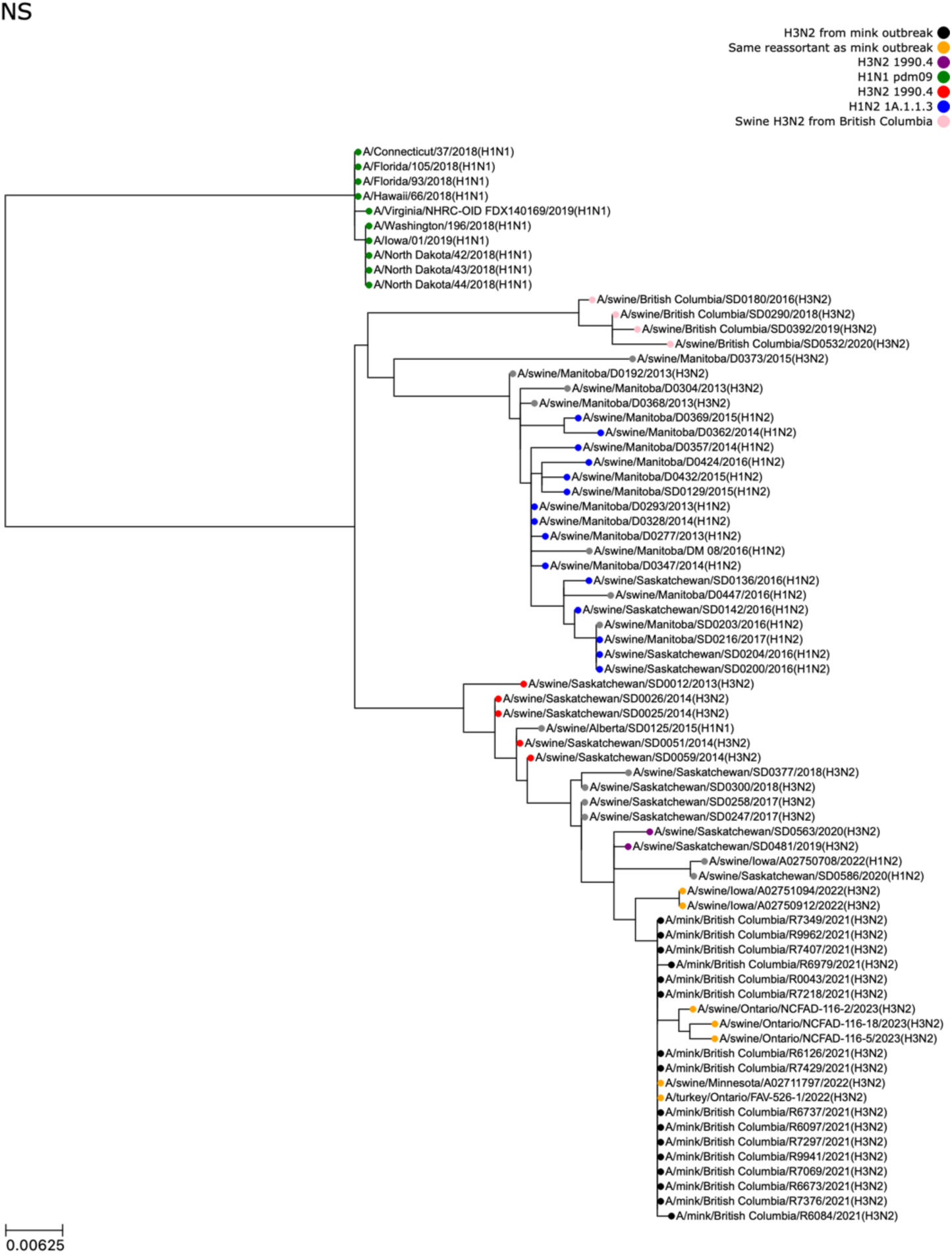
Phylogenetic tree depicting evolutionary relationships between NS segments from the influenza A virus (IAV) detected in mink outbreak and other closely-related IAVs. Other IAV sequences were obtained from NCBI or the Canadian Food Inspection Agency. NS segments from reference viruses were included in the tree if any segment from those viruses was one of the top matches for any segment of the mink virus. The phylogeny indicates the IAV detected in the mink outbreak (black leaves) obtained its NS segment from a swine H3N2 clade 1990.4 virus (purple leaves), which itself obtained this NS segment from an earlier H3N2 clade 1990.4 virus (red leaves).

**Table S1:**
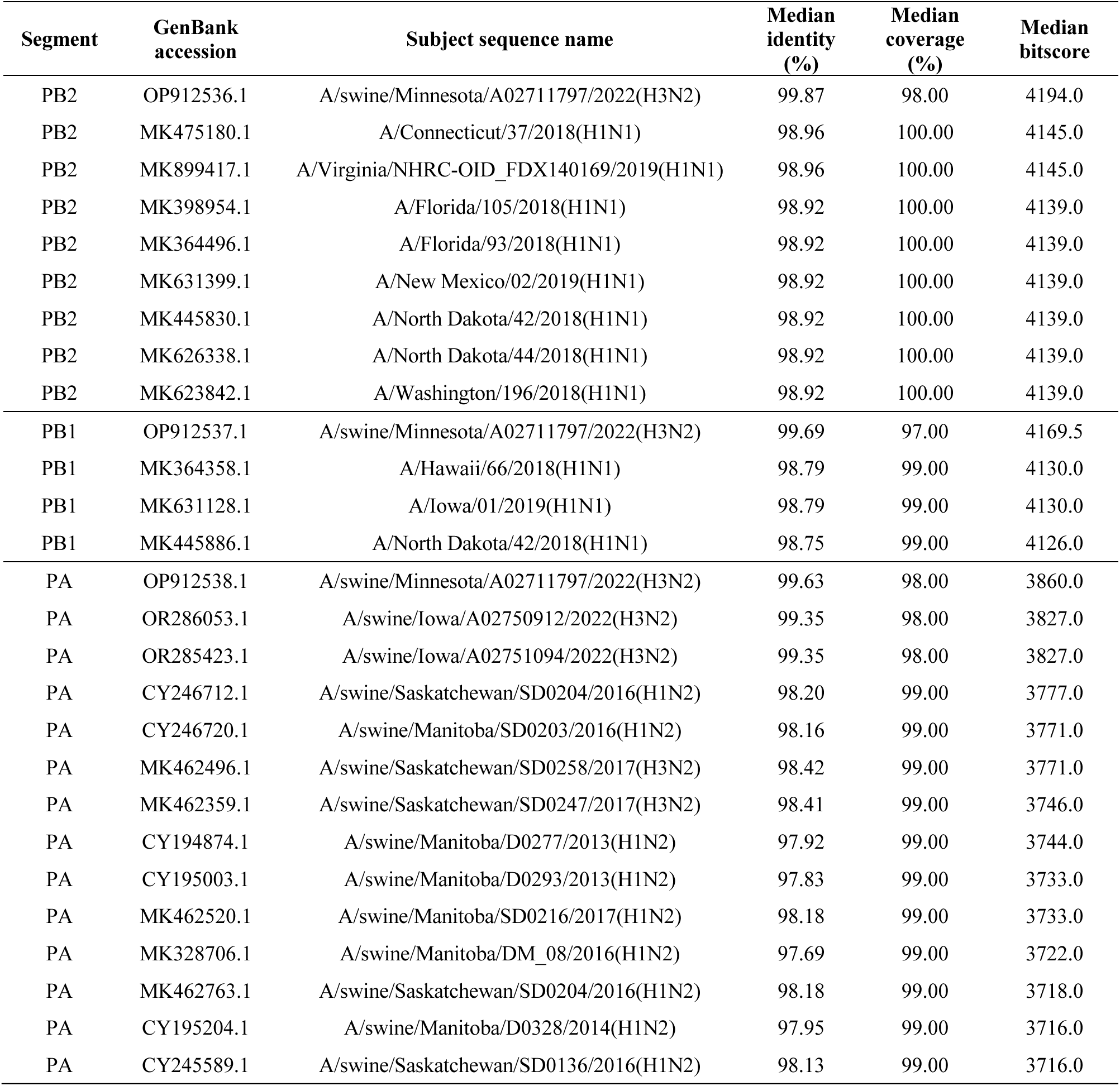
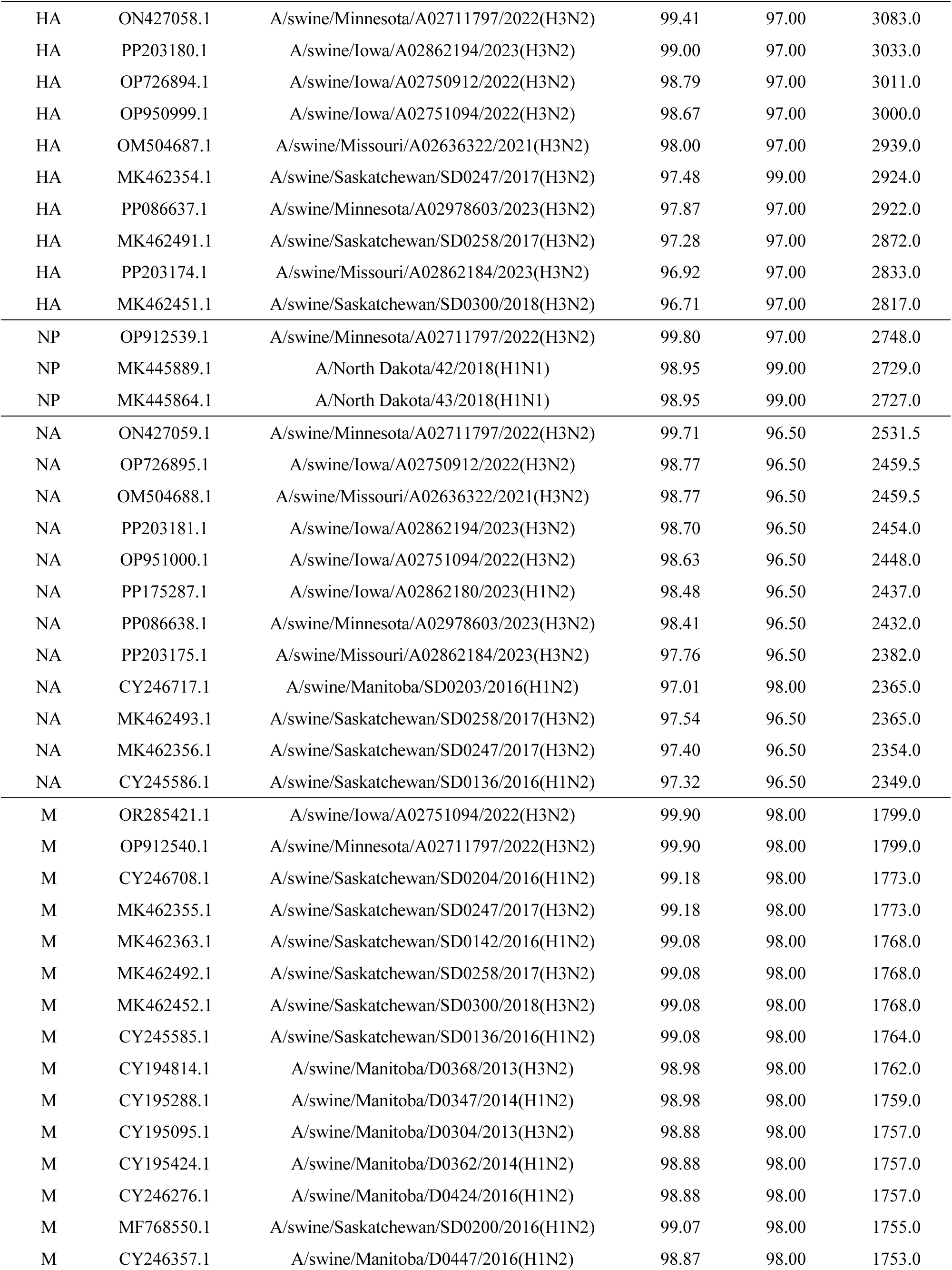
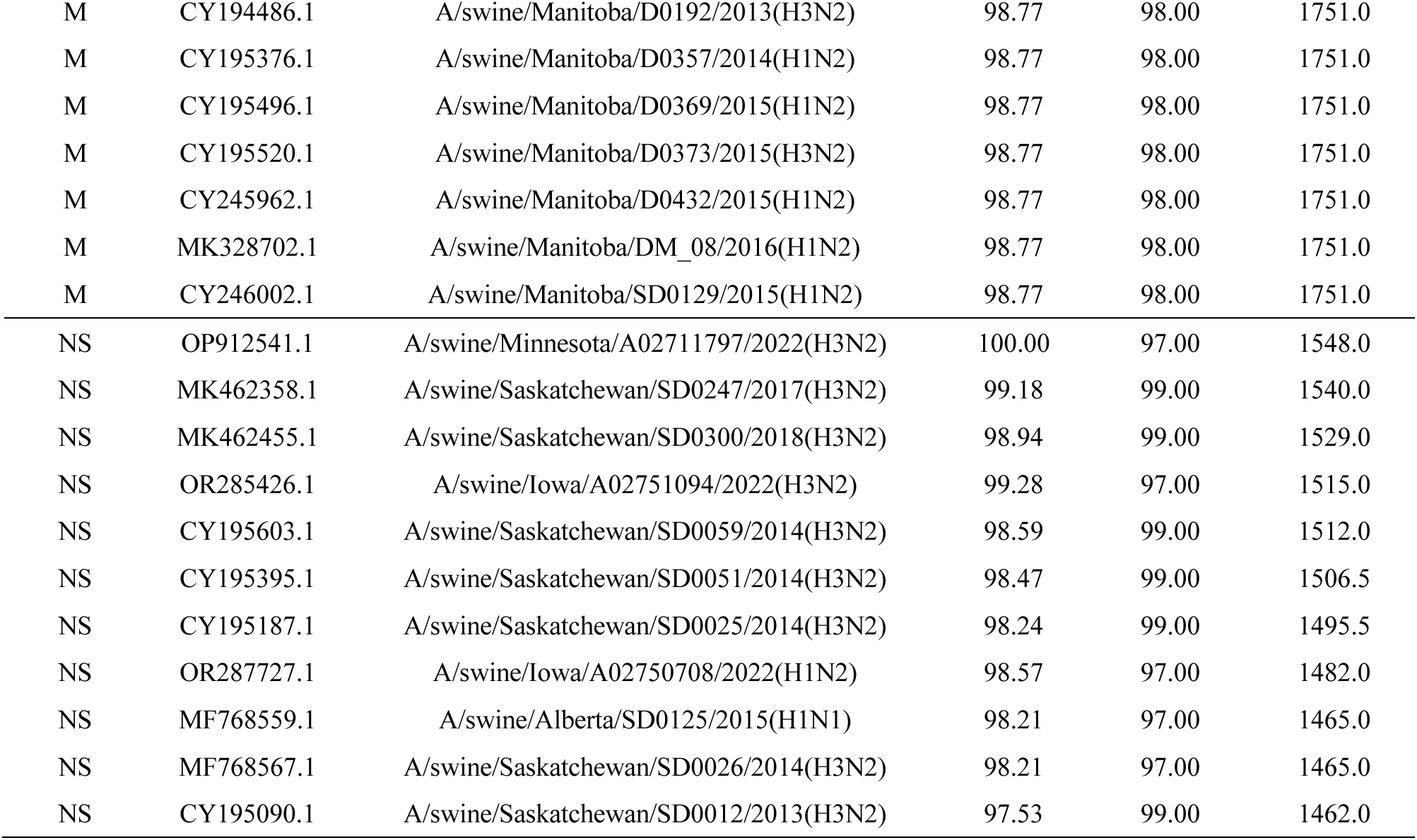
Best matches in GenBank for H3N2 influenza A virus segments detected in farmed mink in British Columbia, Canada. Alignments were performed using blastn against the NCBI nucleotide collection on Feb 19, 2024. For each reference sequence, the median nucleotide identity, subject coverage, and bitscore was calculated from all mink segment sequences that aligned to it. Results were limited to the top *n* subject sequences based on median bitscore (*n*=3 for PB2, PB1, and NP and *n*=10 for other segments).

## Notes

### Competing Interest Statement

The authors have declared no competing interest.

## References

1. Zhao C, Pu J. Influence of Host Sialic Acid Receptors Structure on the Host Specificity of Influenza Viruses. Viruses. 2022;14(10):2141. doi:10.3390/v14102141

2. Zhao P, Sun L, Xiong J, et al. Semiaquatic mammals might be intermediate hosts to spread avian influenza viruses from avian to human. Sci Rep. 2019;9:11641. doi:10.1038/s41598-019-48255-5

3. Ng PSK, Böhm R, Hartley-Tassell LE, et al. Ferrets exclusively synthesize Neu5Ac and express naturally humanized influenza A virus receptors. Nat Commun. 2014;5:5750. doi:10.1038/ncomms6750

4. Ma W, Lager KM, Vincent AL, Janke BH, Gramer MR, Richt JA. The Role of Swine in the Generation of Novel Influenza Viruses. Zoonoses and Public Health. 2009;56(6-7):326–337. doi:10.1111/j.1863-2378.2008.01217.x

5. Nelli RK, Kuchipudi SV, White GA, Perez BB, Dunham SP, Chang KC. Comparative distribution of human and avian type sialic acid influenza receptors in the pig. BMC Veterinary Research. 2010;6(1):4. doi:10.1186/1746-6148-6-4

6. Ma W, Kahn RE, Richt JA. The pig as a mixing vessel for influenza viruses: Human and veterinary implications. J Mol Genet Med. 2008;3(1):158–166.

7. Abdelwhab EM, Mettenleiter TC. Zoonotic Animal Influenza Virus and Potential Mixing Vessel Hosts. Viruses. 2023;15(4):980. doi:10.3390/v15040980

8. Yagyu K, Yanagawa R, Matsuura Y, Fukushi H, Kida H, Noda H. Serological survey of influenza A virus infection in mink. Nihon Juigaku Zasshi. 1982;44(4):691–693. doi:10.1292/jvms1939.44.691

9. Gagnon CA, Spearman G, Hamel A, et al. Characterization of a Canadian Mink H3N2 Influenza A Virus Isolate Genetically Related to Triple Reassortant Swine Influenza Virus. J Clin Microbiol. 2009;47(3):796–799. doi:10.1128/JCM.01228-08

10. Clayton MJ, Kelly EJ, Mainenti M, et al. Pandemic lineage 2009 H1N1 influenza A virus infection in farmed mink in Utah. J VET Diagn Invest. 2022;34(1):82-85. doi:10.1177/10406387211052966

11. Agüero M, Monne I, Sánchez A, et al. Highly pathogenic avian influenza A(H5N1) virus infection in farmed minks, Spain, October 2022. Euro Surveill. 2023;28(3):2300001. doi:10.2807/1560-7917.ES.2023.28.3.2300001

12. Pomorska-Mól M, Włodarek J, Gogulski M, Rybska M. Review: SARS-CoV-2 infection in farmed minks – an overview of current knowledge on occurrence, disease and epidemiology. Animal. 2021;15(7):100272. doi:10.1016/j.animal.2021.100272

13. Koopmans M. SARS-CoV-2 and the human-animal interface: outbreaks on mink farms. Lancet Infect Dis. 2021;21(1):18–19. doi:10.1016/S1473-3099(20)30912-9

14. Liu J, Li Z, Cui Y, Yang H, Shan H, Zhang C. Emergence of an Eurasian avian-like swine influenza A (H1N1) virus from mink in China. Vet Microbiol. 2020;240:108509. doi:10.1016/j.vetmic.2019.108509

15. Bo-Shun Z, Li LJ, Qian Z, et al. Co-infection of H9N2 influenza virus and Pseudomonas aeruginosa contributes to the development of hemorrhagic pneumonia in mink. Vet Microbiol. 2020;240:108542. doi:10.1016/j.vetmic.2019.108542

16. Jiang W, Wang S, Zhang C, et al. Characterization of H5N1 highly pathogenic mink influenza viruses in eastern China. Vet Microbiol. 2017;201:225–230. doi:10.1016/j.vetmic.2017.01.028

17. Peng L, Chen C, Kai-yi H, et al. Molecular characterization of H9N2 influenza virus isolated from mink and its pathogenesis in mink. Vet Microbiol. 2015;176(1-2):88–96. doi:10.1016/j.vetmic.2015.01.009

18. Yoon KJ, Schwartz K, Sun D, Zhang J, Hildebrandt H. Naturally occurring Influenza A virus subtype H1N2 infection in a Midwest United States mink (Mustela vison) ranch. J Vet Diagn Invest. 2012;24(2):388–391. doi:10.1177/1040638711428349

19. Hempel EM, Bharmal A, Li G, et al. Prospective, clinical comparison of self-collected throat-bilateral nares swabs and saline gargle compared to health care provider collected nasopharyngeal swabs among symptomatic outpatients with potential SARS-CoV-2 infection. J Assoc Med Microbiol Infect Dis Can. 8(4):283–298. doi:10.3138/jammi-2023-0002

20. Zhou B, Donnelly ME, Scholes DT, et al. Single-Reaction Genomic Amplification Accelerates Sequencing and Vaccine Production for Classical and Swine Origin Human Influenza A Viruses. J Virol. 2009;83(19):10309–10313. doi:10.1128/JVI.01109-09

21. Hickman R, Nguyen J, Lee TD, et al. Rapid, High-Throughput, Cost Effective Whole Genome Sequencing of SARS-CoV-2 Using a Condensed One Hour Library Preparation of the Illumina DNA Prep Kit. Published online February 8, 2022:2022.02.07.22269672. doi:10.1101/2022.02.07.22269672

22. Camacho C, Coulouris G, Avagyan V, et al. BLAST+: architecture and applications. BMC Bioinformatics. 2009;10:421. doi:10.1186/1471-2105-10-421

23. Guindon S, Lethiec F, Duroux P, Gascuel O. PHYML Online—a web server for fast maximum likelihood-based phylogenetic inference. Nucleic Acids Res. 2005;33(Web Server issue):W557-W559. doi:10.1093/nar/gki352

24. Edgar RC. MUSCLE: multiple sequence alignment with high accuracy and high throughput. Nucleic Acids Res. 2004;32(5):1792–1797. doi:10.1093/nar/gkh340

25. Olsen CW, Karasin AI, Carman S, et al. Triple Reassortant H3N2 Influenza A Viruses, Canada, 2005 - Volume 12, Number 7—July 2006 - Emerging Infectious Diseases journal - CDC. doi:10.3201/eid1207.060268

26. Anderson TK, Nelson MI, Kitikoon P, Swenson SL, Korslund JA, Vincent AL. Population dynamics of cocirculating swine influenza A viruses in the United States from 2009 to 2012. Influenza Other Respir Viruses. 2013;7(Suppl 4):42–51. doi:10.1111/irv.12193

27. Walia RR, Anderson TK, Vincent AL. Regional patterns of genetic diversity in swine influenza A viruses in the United States from 2010 to 2016. Influenza and Other Respiratory Viruses. 2019;13(3):262. doi:10.1111/irv.12559

28. Choi YK, Lee JH, Erickson G, et al. H3N2 influenza virus transmission from swine to turkeys, United States. Emerg Infect Dis. 2004;10(12):2156–2160. doi:10.3201/eid1012.040581

29. Kuchinski KS, Coombe M, Mansour SC, et al. Targeted genomic sequencing of avian influenza viruses in wetland sediment from wild bird habitats. Appl Environ Microbiol. 2024;90(2):e0084223. doi:10.1128/aem.00842-23

30. Nituch LA, Bowman J, Wilson PJ, Schulte-Hostedde AI. Aleutian mink disease virus in striped skunks (Mephitis mephitis): evidence for cross-species spillover. J Wildl Dis. 2015;51(2):389–400. doi:10.7589/2014-05-141

31. Gillett NP, Cannon AJ, Malinina E, et al. Human influence on the 2021 British Columbia floods. Weather and Climate Extremes. 2022;36:100441. doi:10.1016/j.wace.2022.100441

32. Ramey AM, Reeves AB, Lagassé BJ, et al. Evidence for interannual persistence of infectious influenza A viruses in Alaska wetlands. Science of The Total Environment. 2022;803:150078. doi:10.1016/j.scitotenv.2021.150078

33. Ramey AM, Reeves AB, Drexler JZ, et al. Influenza A viruses remain infectious for more than seven months in northern wetlands of North America. Proceedings of the Royal Society B: Biological Sciences. 2020;287(1934):20201680. doi:10.1098/rspb.2020.1680

34. Karasin AI, Olsen CW, Brown IH, Carman S, Stalker M, Josephson G. H4N6 influenza virus isolated from pigs in Ontario. Can Vet J. 2000;41(12):938–939.

35. Corzo CA, Culhane M, Dee S, Morrison RB, Torremorell M. Airborne Detection and Quantification of Swine Influenza A Virus in Air Samples Collected Inside, Outside and Downwind from Swine Barns. PLOS ONE. 2013;8(8):e71444. doi:10.1371/journal.pone.0071444

36. Scoizec A, Niqueux E, Thomas R, Daniel P, Schmitz A, Le Bouquin S. Airborne Detection of H5N8 Highly Pathogenic Avian Influenza Virus Genome in Poultry Farms, France. Frontiers in Veterinary Science. 2018;5. Accessed September 19, 2023. https://www.frontiersin.org/articles/10.3389/fvets.2018.00015

37. Ypma RJF, Jonges M, Bataille A, et al. Genetic data provide evidence for wind-mediated transmission of highly pathogenic avian influenza. J Infect Dis. 2013;207(5):730–735. doi:10.1093/infdis/jis757

38. Ssematimba A, Hagenaars TJ, de Jong MCM. Modelling the Wind-Borne Spread of Highly Pathogenic Avian Influenza Virus between Farms. PLoS One. 2012;7(2):e31114. doi:10.1371/journal.pone.0031114

39. Jonges M, van Leuken J, Wouters I, Koch G, Meijer A, Koopmans M. Wind-Mediated Spread of Low-Pathogenic Avian Influenza Virus into the Environment during Outbreaks at Commercial Poultry Farms. PLoS One. 2015;10(5):e0125401. doi:10.1371/journal.pone.0125401

40. Okazaki K, Yanagawa R, Kida H, Noda H. Human influenza virus infection in mink: Serological evidence of infection in summer and autumn. Veterinary Microbiology. 1983;8(3):251–257. doi:10.1016/0378-1135(83)90077-9

41. Sun H, Li F, Liu Q, et al. Mink is a highly susceptible host species to circulating human and avian influenza viruses. Emerg Microbes Infect. 10(1):472–480. doi:10.1080/22221751.2021.1899058

42. Zhang C, Xuan Y, Shan H, et al. Avian influenza virus H9N2 infections in farmed minks. Virol J. 2015;12:180. doi:10.1186/s12985-015-0411-4

43. Åkerstedt J, Valheim M, Germundsson A, et al. Pneumonia caused by influenza A H1N1 2009 virus in farmed American mink (Neovison vison). Veterinary Record. 2012;170(14):362–362. doi:10.1136/vr.100512

44. Mok CKP, Qin K. Mink infection with influenza A viruses: an ignored intermediate host? One Health Advances. 2023;1(1):5. doi:10.1186/s44280-023-00004-0

